## Supplementary Figures 1-7 for "Translaminar Recurrence from Layer 5 Suppresses Superficial Cortical Layers"

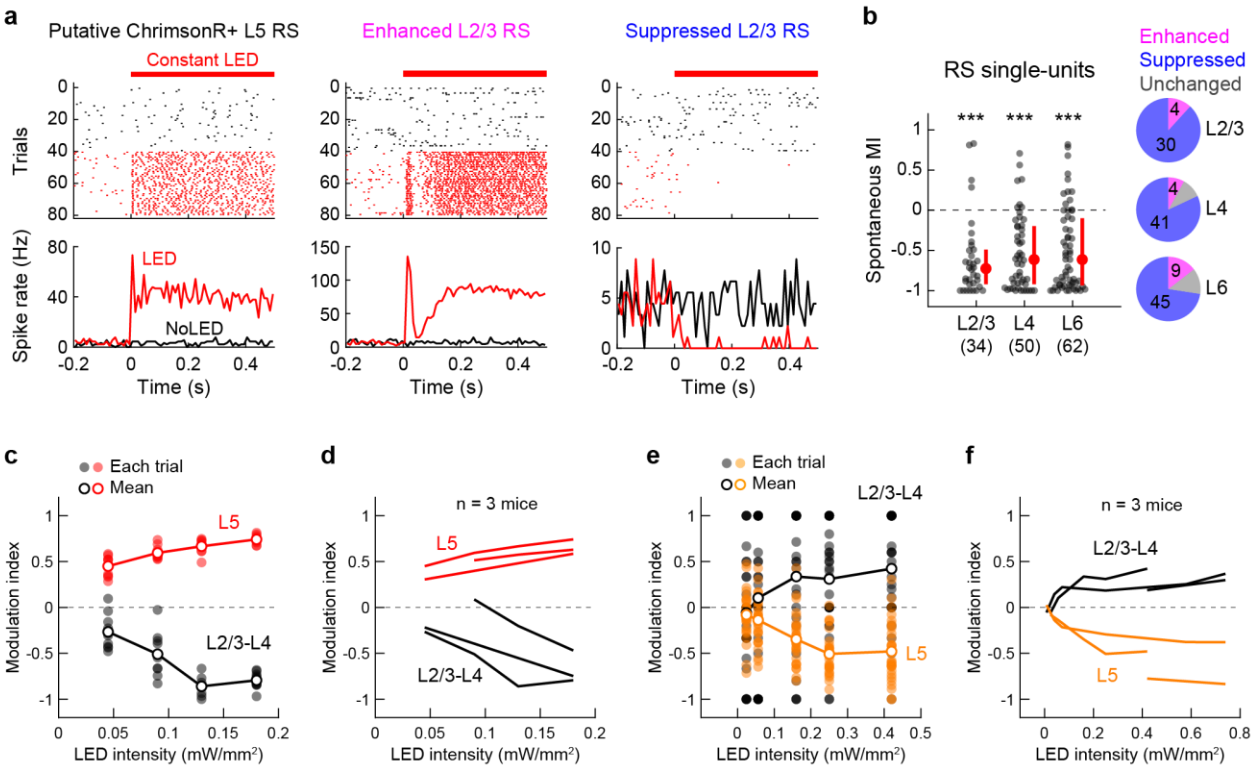


**Supplementary Figure 1. LED profiles do not affect the impacts of L5 optogenetic manipulations on cortical activity. (a,b)** Data for ChrimsonR-mediated L5 activation using constant-intensity LED without an onset ramp. **(a)** Rasters and PSTHs of spontaneous firing with (red) and without (black) LED in a representative photoactivated L5 RS single-unit (left), an enhanced L2/3 RS single-unit (middle), and a suppressed L2/3 RS single-unit (right). Red bars, 625 nm illumination. **(b)** Left, scatter plots showing MI for the spontaneous firing of RS single-units in each layer across mice (9 mice; n = 34, 50, 62 units for L2/3, L4, L6). Red dots and bars represent median and 25th and 75th percentiles. Right, pie charts showing the fraction of significantly enhanced (pink), suppressed (blue), and unchanged (gray) units. ***$p < 0.001$. Wilcoxon signed rank test with Bonferroni correction. **(c,d)** Data for ChrimsonR-mediated L5 activation with low LED intensities. **(c)** MI of L2/3–L4 (black) and L5 (red) RS multi-unit spontaneous activity across LED intensities in a representative mouse. Shaded circles show individual trials, and open circles indicate the mean. **(d)** MI change of individual mice (n = 3 mice). There is a monotonous increase in the optogenetic effect size with LED intensity. **(e,f)** Data for eNpHR3-mediated L5 inactivation with low LED intensities. **(e)** MI of L2/3–L4 (black) and L5 (amber) RS multi-unit spontaneous activity across LED intensities in a representative mouse. **(f)** MI change of individual mice (n = 3 mice). There is a monotonous increase in the optogenetic effect size with LED intensity.

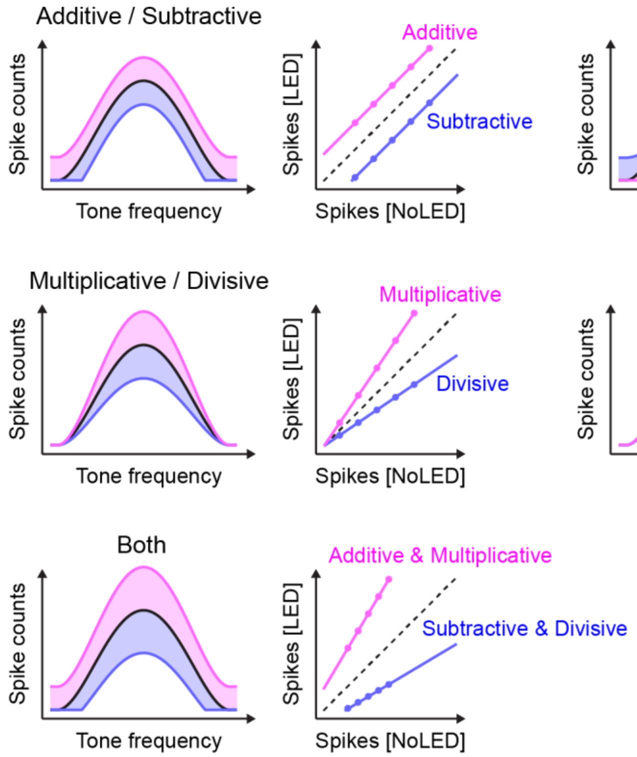


**Supplementary Figure 2. Classification of five types of gain-control transformations.** Schematics illustrating our five classifications of gain control transformations in Fig. 3 and Supplementary Fig. 6. In each type, the left panel shows frequency tuning curves with (pink and blue) and without (black) optogenetic manipulation. The right panel shows tone-evoked spike counts in LED and No LED conditions. Solid lines show threshold-linear fit, and dotted lines indicate unity.

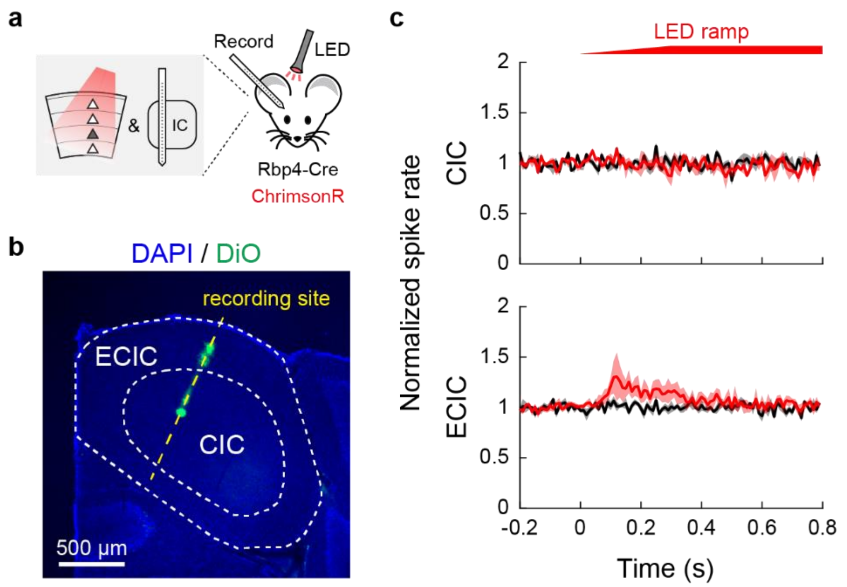


**Supplementary Figure 3. L5 activation does not suppress the inferior colliculus. (a)** Schematics illustrating the recording of the inferior colliculus with optogenetic activation of A1 L5 neurons. **(b)** Coronal section of the inferior colliculus in a representative mouse. Recording site (yellow dotted line) was identified by DiO signal. CIC: central nucleus of the inferior colliculus. ECIC: external cortex of the inferior colliculus. **(c)** Normalized PSTHs of multi-unit activity combining spikes in CIC (top) and ECIC (bottom) with (red) and without (black) L5 activation. The traces are the average of 5 recordings (3 mice), and the shading shows SEM. Red bar, 625 nm LED.

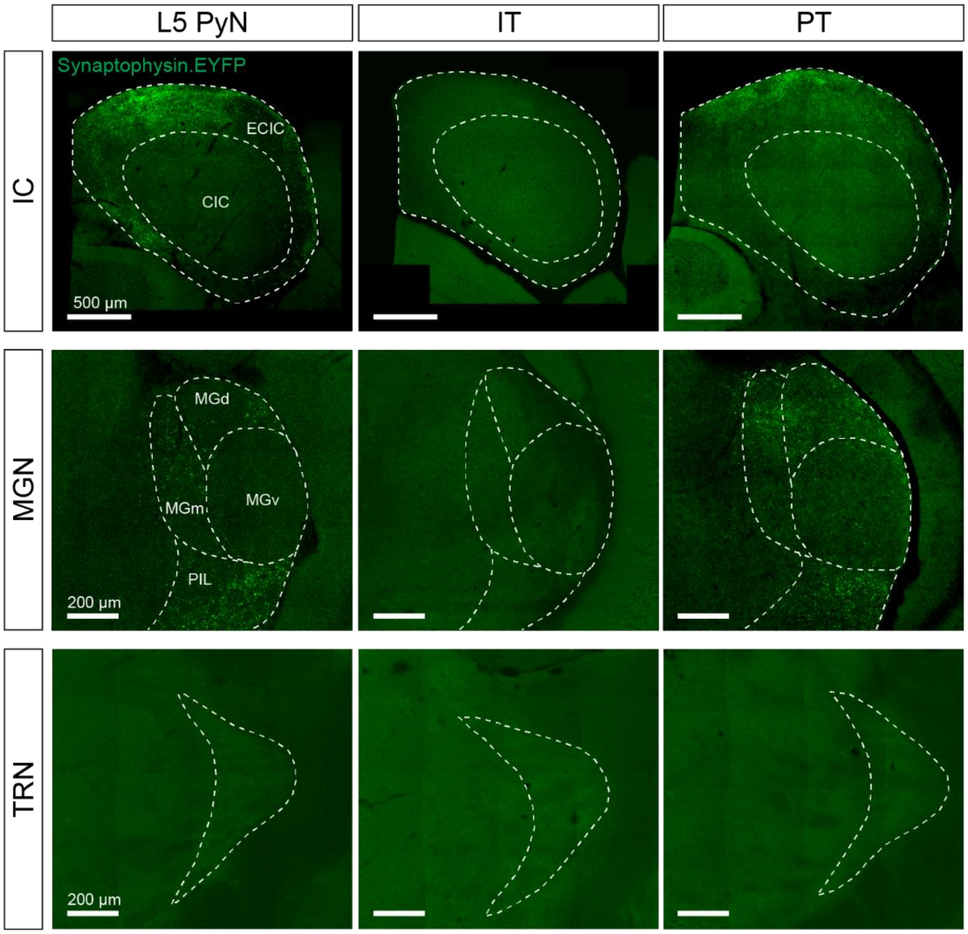


**Supplementary Figure 4. PT but not IT neurons project to the thalamus and inferior colliculus.** Coronal sections of the inferior colliculus (IC; top row), medial geniculate nucleus (MGN; middle row), and thalamic reticular nucleus (TRN; bottom row), from mice expressing synaptophysin.EYFP in distinct L5 pyramidal neuron populations. AAV8.2.hEF1a.DIO.synaptophysin.EYFP was injected into A1 of Rbp4-Cre mice (pan-L5 pyramidal cells; left), Tlx3-Cre mice (IT neurons; middle), and mice with CAV2-Cre injections in the ECIC (PT neurons; right). Pan-L5 pyramidal cells and PT neurons projected to the non-primary subdivisions of the IC (ECIC) and MGN (MGd, MGm, and PIL) while IT neurons did not. No projections were observed in TRN. Dotted lines indicate the boundaries of the regions. ECIC: external cortex of IC. CIC: central nucleus of IC. MGv: ventral part of MGN. MGd: dorsal part of MGN. MGm: medial part of MGN. PIL: posterior intralaminar thalamic nucleus.

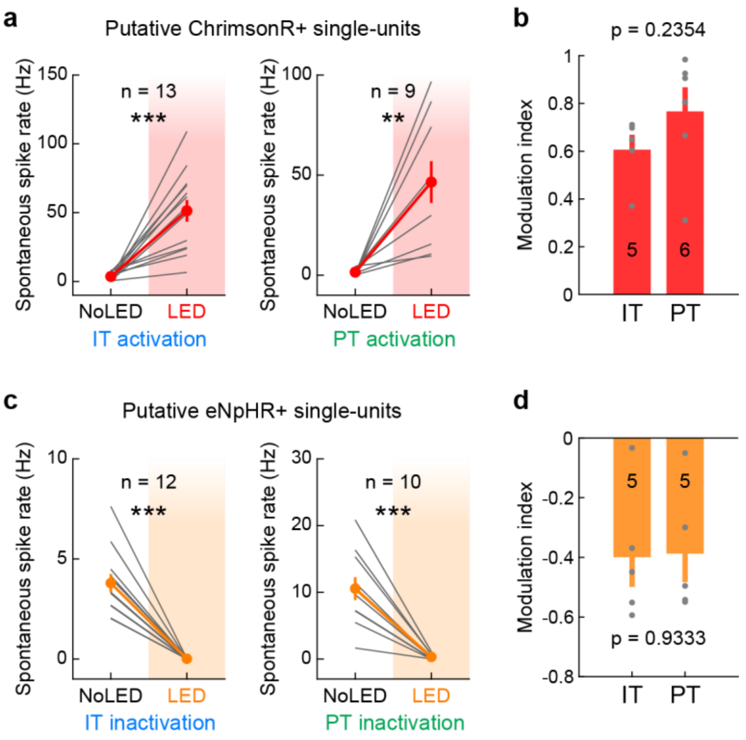


**Supplementary Figure 5. Optogenetic modulation magnitudes are comparable between IT and PT manipulations. (a)** Spontaneous firing rates of putative ChrimsonR-expressing L5 single-units with and without LED. Left, IT activation (n = 5 mice, 13 units). Right, PT activation (n = 6 mice, 9 units). Gray lines show individual units, and red lines indicate mean ± SEM. $^{**}p < 0.01$, $^{***}p < 0.001$. Two-sided paired *t*-test. **(b)** Comparison of MI for spontaneous RS multi-unit activity in photoactivated L5 depth bins between IT and PT activations. Data are mean ± SEM, overlaid with individual mice (n = 5, 6 mice for IT, PT). p = 0.2354, two-sample *t*-test. **(c)** Spontaneous firing rates of putative eNpHR3-expressingL5 single-units with and without LED. Left, IT inactivation (n = 5 mice, 12 units). Right, PT inactivation (n = 5 mice, 10 units). Note that the spontaneous firing rate in No LED condition is high in these data since units with low basal firing rates failed to be detected as putative eNpHR3-expressing units. **(d)** Comparison of MI for spontaneous RS multi-unit activity in photosuppressed L5 depth bins between IT and PT inactivations (n = 5, 5 mice for IT, PT). p = 0.9333.

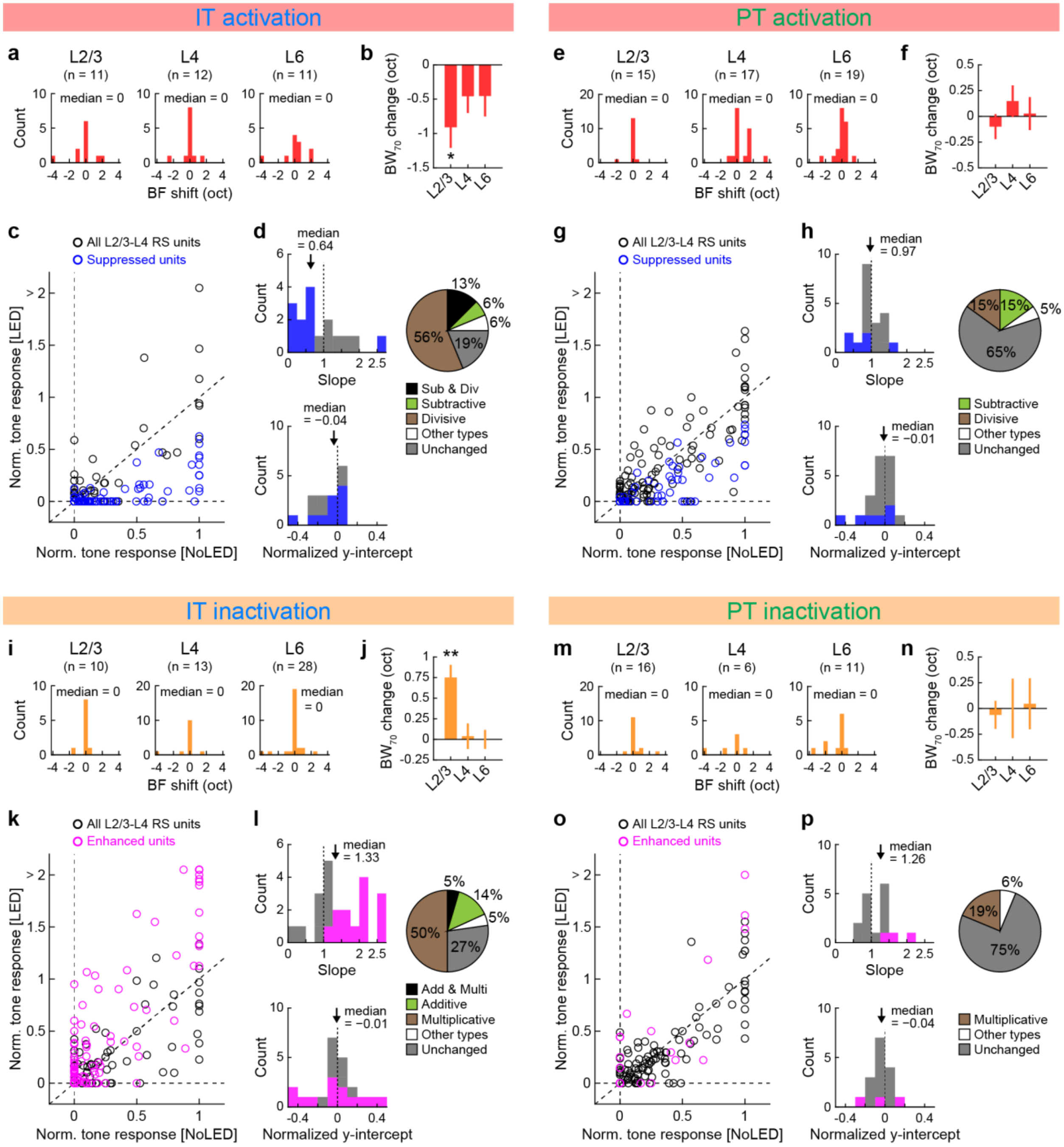


**Supplementary Figure 6. IT neurons modulate tone responses in the superficial layers more robustly than PT neurons. (a–d)** Data for A1 tone-evoked responses during IT neuron activation with ChrimsonR. **(a)** Distribution of the best frequency (BF) shift from No LED to LED conditions (n = 11, 12, 11 tone-responsive units for L2/3, L4, L6). **(b)** LED-induced changes in the population frequency tuning bandwidth at 70 dB SPL ($BW_{70}$) in each layer. *p =

0.038, one-sample *t*-test with Bonferroni correction. **(c)** Summary data showing well-fit RS single-units in L2/3–L4 (n = 16 single-units, 144 unit-tone pairs). Spike counts are normalized to the BF response without LED in each unit. Blue circles show suppressed units. Suppressed units (blue circles) were defined by the MI of the total spike counts for all responsive frequencies; therefore, some of the data points for suppressed units could appear above the unity line. **(d)** Distribution of best-fit slope (top left) and normalized y-intercept (bottom left) for all L2/3–L4 RS single-units that were fit well with threshold-linear functions. Blue bars indicate units with significant suppression of tone-evoked activity. Arrows show median. Right, a pie chart showing the fraction of units with divisive (brown), subtractive (green), both suppression (black), other modulations (white), and no change (gray) (n = 16 units). **(e–h)** Same as in (a–d) but for PT neuron activation with ChrimsonR. **(e,f)** n = 15, 17, 19 tone-responsive units for L2/3, L4, L6. **(g,h)** n = 20 single-units, 180 unit-tone pairs. **(i–l)** Same as in (a–d) but for IT inactivation with eNpHR3. **(i,j)** n = 10, 13, 28 tone-responsive units for L2/3, L4, L6. **p = 0.0026. **(k–l)** n = 22 single-units, 198 unit-tone pairs. Pink circles and bars indicate units with significant suppression of tone-evoked activity. **(m–p)** Same as in (a–d) but for PT inactivation with eNpHR3. **(m,n)** n = 16, 6, 11 tone-responsive units for L2/3, L4, L6. **(o,p)** n = 16 single-units, 144 unit-tone pairs.

Taken together, IT neuron manipulations changed the frequency tuning of superficial layers similarly to pan-L5 manipulations (Fig. 3), whereas PT neuron manipulations did not alter tone processing significantly.

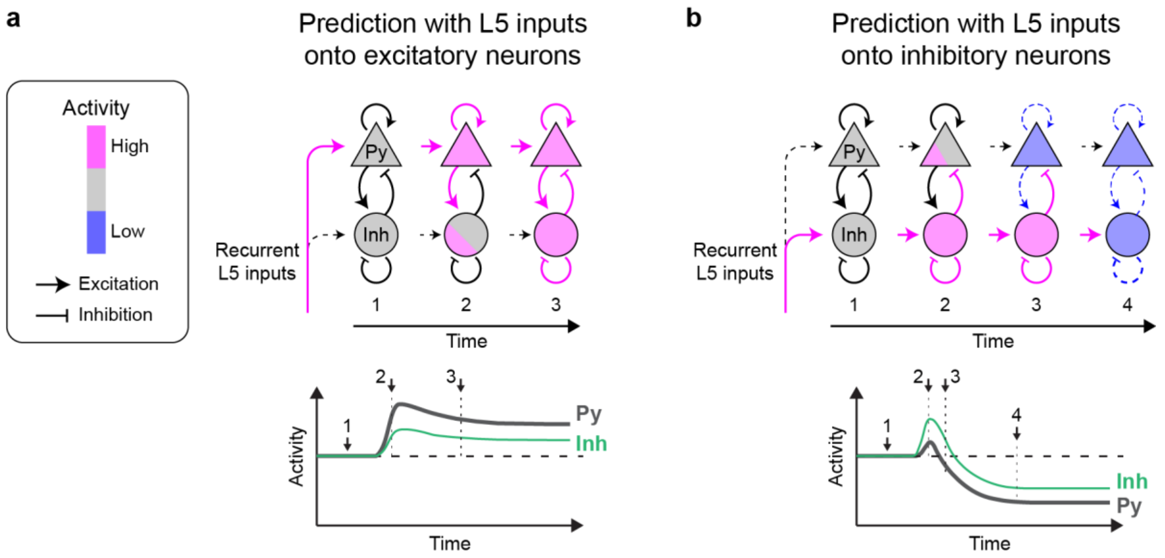


**Supplementary Figure 7. Models of translaminar L5 recurrence with the inhibition-stabilized network. (a)** Cartoons summarizing the predicted dynamics of L2/3 cortical activity when L5 excitatory neurons synapse predominantly onto pyramidal cells. Top, diagram of the temporal dynamics of inhibitory-stabilized networks (ISN), adapted from Ozeki et al[4]. The network is composed of two cell populations: excitatory pyramidal cells (triangles) and inhibitory neurons (circles). Pink and blue colors represent increased and decreased activity levels, respectively. Numbers indicate the corresponding time points in the bottom panel. Bottom, time course of the activity of pyramidal cells (Py; black) and inhibitory neurons (Inh; green). Excitatory inputs onto pyramidal cells increase the activity of both pyramidal cells and inhibitory neurons. **(b)** Prediction when L5 excitatory neurons synapse predominantly onto L2/3 inhibitory neurons. Excitatory inputs transiently increase the activity of inhibitory neurons (2). However, the recruited inhibitory neurons suppress pyramidal cells and reduce local recurrent excitation within the L2/3 network (3). Since cortical circuits are dominated by local recurrent excitation in the ISN model, the resulting reduction in the excitatory input onto inhibitory neurons overcomes the initial excitation from L5, resulting in a paradoxical reduction of inhibitory neurons firing (4)[4,6,7,10,46].
